# A flexible implant towards acute intrapancreatic electrophysiology

**DOI:** 10.1101/2023.03.17.532901

**Authors:** Domenic Pascual, Lisa Brauns, Ruth Domes, Matthias Tisler, Marco Kögel, Angelika Stumpf, Andreas Kirschniak, Jens Rolinger, Udo Kraushaar, Peter D. Jones

**Affiliations:** NMI Natural and Medical Sciences Institute at the University of Tübingen, Reutlingen, Germany; University Hospital Tübingen, Tübingen, Germany; Maria Hilf Hospital, Mönchengladbach, Germany

**Keywords:** depth recording, pancreas, beta cells, diabetes mellitus, islets of Langerhans, microelectrode array

## Abstract

Microelectrode arrays (MEAs) have proven to be a powerful tool to study electrophysiological processes over the last decades with most technology developed for investigation of the heart or brain. Other targets in the field of bioelectronic medicine are the peripheral nervous system and its innervation of various organs. Beyond the heart and nervous systems, the beta cells of the pancreatic islets of Langerhans generate action potentials during the production of insulin. In vitro experiments have demonstrated that their activity is a biomarker for blood glucose levels, suggesting that recording their activity in vivo could support patients suffering from diabetes mellitus with long-term automated read-out of blood glucose concentrations.

Here, we present a flexible polymer-based implant having 64 low impedance microelectrodes designed to be implanted to a depth of 10 mm into the pancreas. As a first step, the implant will be used in acute experiments in pigs to explore the electrophysiological processes of the pancreas in vivo. Beyond use in the pancreas, our flexible implant and simple implantation method may also be used in other organs such as the brain.

## 1 Introduction

Electrodes implanted into the human brain or heart have shown impressive clinical results for decades. In pacemakers and implanted defibrillators, the electrodes record cardiac signals and can artificially stimulate an absent heartbeat (Trohman, Kim and Pinski, 2004) or even end periods of fibrillation by delivering an electric shock (DiMarco, 2003). Other clinical targets of bioelectronic medicine include the brain and central nervous system, the peripheral nervous system (including autonomic nerves), muscles, and organs with other electrogenic cells (Long *et al*., 2021). Specifically, the beta cells of the pancreas are electrically active during insulin release. Previously, we have demonstrated the measurement of such electrical activity from isolated pancreatic islets of Langerhans and its correlation with glucose concentration (Pfeiffer *et al*., 2011; Schönecker *et al*., 2015).

In Europe more than 56 million people currently suffer from diabetes mellitus, with a rising prevalence (Tamayo *et al*., 2014). The International Diabetes Federation estimates over 536 million patients worldwide (International Diabetes Federation, 2021). Diabetes mellitus is predominantly a metabolic disorder of the pancreas characterized by an increase in blood glucose levels. In the course of the disease, the pancreatic beta cells lose their ability to produce insulin, which leads to hyperglycaemia in affected patients and further to insulin resistance and thus to the manifestation of diabetes mellitus (Inzucchi *et al*., 2012). Type 1 diabetes mellitus is an autoimmune disease leading to the destruction of insulin-producing beta cells and causing absolute insulin deficiency, while type 2 diabetes mellitus is a metabolic disorder characterized by both insulin resistance and inadequate insulin production. Affected patients must independently monitor their blood glucose levels and respond to them (Pfeiffer and Klein, 2014). Hyperglycaemia in untreated diabetes mellitus or in patients with low compliance causes microvascular changes which manifest in comorbidities such as diabetic polyneuropathy, cardiovascular disease, and increased risk of cancer (Tsuchitani, Sato and Kokoshima, 2016).

Therefore, we see the need for new therapeutic options that reduce the patients’ burden of glucose monitoring in case of diabetes mellistus type 2, thereby improving compliance and improving patient health. We envision a future closed-loop device for long-term monitoring of the electrical activity of the pancreas coupled to a system for insulin release.

The human pancreas is 12–15 cm long and its neck is about 2 cm wide. The porcine pancreas is a useful model due to its similar size and compact anatomy (Standring, 2016). Smaller animals such as rats have a smaller pancreas which is in part dispersed within the mesentery (Ionescu-Tirgoviste *et al*., 2015; Tsuchitani, Sato and Kokoshima, 2016). The pancreas consists of 90 % exocrine tissue and 4.5 % endocrine tissue. The endocrine tissue consists of islets of Langerhans, which are spherically organized cell assemblies containing mostly beta cells, which are responsible for the production of insulin. A human pancreas contains a total of 3.2 to 14.8 million of these islets. The average diameter of a pancreatic islet is about 110 µm (Ionescu-Tirgoviste *et al*., 2015). Exocrine tissue of the pancreas produces digestive enzymes that can be released upon injury, causing pancreatitis. Therefore, potential tissue damage must be reduced by the design specifications of the implant.

The electrical activity of beta cells concurrent with insulin release was first confirmed in 1968 in ex vivo mouse pancreas (Dean and Matthews, 1968) and later measured in vivo in the mouse (Sánchez-Andrés, Gomis and Valdeolmillos, 1995); both studies used glass microelectrodes to measure the intracellular potential. Intracellular recordings have a short duration (minutes or hours). In vitro experiments using planar glass-based microelectrode arrays allow electrical recording of islets at defined glucose and drug concentrations (Schönecker *et al*., 2014), including using human islets (Schönecker *et al*., 2015).

A patent from 2001 describes the method of electropancreatography and an apparatus for recording the electrical activity of the pancreas (Harel, Tamar *et al*., 2001). Another patent, in addition to the ability to sense beta cell activity, relates to delivering a stimulation impulse to the pancreas to trigger insulin production (Richard P.M. Houben, 2000). Along with insulin administration, there are attempts to implant xenografts to allow physiological insulin production as needed.

To our knowledge, no in vivo extracellular recording of the glucose-related electrical activity of pancreatic beta cells using microelectrodes has been reported. We are not aware of any microelectrode array (MEA) device that has been implanted into the pancreas. In developing an implant we have therefore applied knowledge of the biology of the pancreas and the implants for neural electrophysiology.

Neither glass pipettes nor in vitro MEAs are suitable for chronic implantation. In designing an implant, important considerations are (1) reducing injury and immune response due to mechanical mismatch, (2) an appropriate surgical implantation method, (3) maturity of materials and fabrication methods and (4) stability in the body for chronic use.

Differences in stiffness and density between the body and an implant lead to micromechanical movement, resulting in an inflammatory response which leads to scarring around the implant and loss of its function (Lin, Tsao and Kuo, 2018). While rigid MEAs such as Utah arrays made of silicon (having a Young’s modulus of ∼170GPa (Cheung, 2007)), have been used clinically in the brain for decades, their use in the pancreas would additionally pose a risk of pancreatitis. Flexible implants can minimize scarring (Yang *et al*., 2019) and have been demonstrated to record stable signals from neurons for months (Jang *et al*., 2021). They are typically produced using thin films of polyimide or parylene with a Young’s modulus in the range of 2.3 – 10 GPa (Weltman, Yoo and Meng, 2016). Another detail for flexible MEAs is the bending stiffness, which can be reduced by perforation of implants such as in mesh MEAs (Lee *et al*., 2019). The use of even softer or stretchable materials such as elastomer (Young’s modulus of 974 kPa) is also promising (Du *et al*., 2017) but less mature. It is worth noting, that, to the best of our knowledge, no data exists regarding the Young’s modulus of the human pancreas (Singh and Chanda, 2021). Consequently, precise matching of implant stiffness with that of the pancreas remains a challenge. For reference, tissue typically has a Young’s modulus below even elastomers, on the order of 5–500 kPa (Singh and Chanda, 2021).

A disadvantage of flexible implants is their challenging implantation. Deep implantation into tissue requires support devices or complex methods (Apollo *et al*., 2020). The implantation is often the most significant factor for injury. In contrast, after healing of the initial injury, flexible or mesh implants often show minimal or no immune response.

Yang *et al*., 2019 developed a MEA that, when stereotactically implanted into the brain of a rodent, did not elicit a significant immune response. They attributed the very good interaction with the tissue to the small dimensions of the electrodes. Furthermore, the flexible negative photoresist SU-8, which supports the electrodes and conductive tracks was designed with a mesh-like structure for high flexibility. A study over 3 months demonstrated stable recording quality. The authors attribute this to the mechanical properties, which largely eliminate micromovements and thus tissue damage.

Takeuchi *et al*., 2005 reported that a fluidic 3D-channel in the polymer of the implant, filled with a biodegradable material led to temporary stiffening of the implant during implantation. Elsewhere highly flexible MEAs could be delivered to the target tissue with a syringe (Liu *et al*., 2015). To adhere an implant to a needle for a short period of time, Du *et al*., 2017 attached a flexible implant to a steel needle with polyethylene glycol (PEG). After implantation, the PEG dissolves in the body’s tissue and after a short residence time, the steel needle could be explanted leaving the MEA inside of the body. Even an attempt to implant a MEA without a shuttle was performed. To stiffen the MEA it was coated it in a bath of PEG. After solidification in ambient air the MEA could be implanted into brain tissue by a micromanipulator (Guan *et al*., 2019).

Flexible MEAs can be produced using reliable microfabrication methods. Polyimide is compatible with all relevant microfabrication methods. In contrast, many polymers or elastomers may be incompatible due to intolerance to solvents, high thermal expansion coefficients leading to misalignment or insufficient thermal properties (e.g. melting or decomposing at necessary temperatures). Finally, stability is important. Polyimide has proven to be reliable for years in clinical implants (Daschner *et al*., 2017) and titanium nitride electrodes can provide good recordings for months (McDonald *et al*., 2023), however the stability of implants in the pancreas requires investigation.

We have designed and fabricated a flexible 64-electrode polymer implant for extracellular recording of the electrical activity of beta cells within the pancreas. We present a shuttle-based method for implantation to a depth of 10 mm. Results include the fabricated devices, their electrochemical characterization, and demonstration of implantation into hydrogel and *ex vivo* pancreas. The next step will be validation in acute experiments in a pig model. As an outlook, the flexible polymer implant could be connected to wireless systems for chronic implantations. The thin-film flexible implant should allow recording of the glucose-related electrical activity of the pancreas, while inducing minimal immune response.

## 2 Methods

### 2.1 Histology

Pancreatic tissue was first treated in a 1x MEFA fixative solution overnight at 4 °C. All materials used for this section were obtained from Merck KGaA. Degreasing of the tissue was performed by 100 % ethanol with an exposure time of at least 24 hours at 4 °C. In preparation for embedding, rehydration of the degreased tissue was performed in phosphate-buffered saline for 3 x 15 minutes at room temperature and equilibration in Vib-Mix (Gelatine-Albumine Mix) for 12 h at 4 °C. Subsequently, the tissue was placed in 2 x 1000 µl Vib-Mix and additionally 25 µl glutaraldehyde (25 %) in a rectangular aluminum mold. The semithin sections of 25 µm were made on a Leica vibratome. Sections were capped using Mowiol and a coverslip. Subsequently, the light microscopic images were acquired.

### 2.2 Design

The implant was designed consisting of a flexible polymer-based microelectrode array (MEA) component connected to a rigid electrode interface board (EIB). Design decisions intended to ensure a high probability of recording the activity of beta cells in the pancreas, maximize flexibility and minimize size, while keeping a good fabrication yield.

### 2.3 Fabrication

Dummy implants having similar structure, but no microelectrodes or traces were fabricated by laser cutting of 12 µm-thick films of polyimide (Du Pont PI 2611) or parylene C on glass wafers using an LPKF Protolaser U with a nominal spot size of 20 µm. Polyimide was deposited in a single spin-coating step. Parylene C was deposited in a parylene coater (Comelec, La Chaux-de-Fonds, Switzerland) using the precursor DPX-C (GALENTIS S.r.l. a Socio Unico). Both were deposited without adhesion promoters to ensure simple removal from carrier wafers.

Flexible polyimide-based components were produced following established methods. A first layer of polyimide (6 µm) was spin-coated on 100 mm silicon carrier wafers. Electrical traces were produced by sputter deposition of Ti/Au/Ti (400 nm), photolithographic patterning of an etch mask, and a combination of dry etching and wet etching (see Figure S1). A second polyimide layer (also 6 µm) was spin-coated. A hard mask of silicon nitride was structured to enable high resolution patterning. Polyimide was etched by oxygen plasma through the full 12 µm thickness but stopped after 6 µm at the metal layer at the electrodes and bond pads. The hard mask was removed by dry etching. Electrodes were patterned with a lift-off process with sputter deposition of TiN. Finally, polyimide components were detached from the carrier wafers.

An electrode interface board (EIB) was designed using KiCad and produced in a 2-layer process with ENEPIG (electroless nickel, electroless palladium, immersion gold) plating (CONTAG AG). The interface between the polyimide component and the EIB was established by rivet bonding (Stieglitz, Beutel and Meyer, 2000; Steins *et al*., 2022). Polyimide components were manually aligned on EIBs. Each channel was bonded with a single gold bump which achieved a resistance below 5 Ω. After bonding, connectors (Omnetics Nano Strip A79024) were soldered to the EIB. We used two 36-channel connectors per implant for a reference electrode, a counter electrode and 64 microelectrodes, in a design compatible with 64-channel RHD headstages (Intan Technologies).

Thereafter, bond and solder contacts were reinforced and encapsulated with epoxy. Underfill of the polyimide using a low viscosity epoxy (Polytec EP 653) was facilitated by holes in the EIB. Encapsulation with UHU plus endfest 300 was applied to protect the EIB in case of contact with liquids during acute implantation.

### 2.4 Impedance measurements

Measurements were made with a Pt mesh counter electrode and silver chloride reference electrode in phosphate-buffered saline with a potentiostat (Gamry 600+) and an electrochemical multiplexer (ECM8 Gamry Instruments).

### 2.5 Implant–shuttle assembly

A stainless-steel needle with a diameter of 250 µm and a length of 3.5 cm serves as the implantation shuttle, and the flexible shank was attached to the shuttle by polyethylene glycol (9000–11250 g/mol, Carl Roth GmbH + Co. KG). First, the shuttle was dip-coated in liquid PEG at ∼75 °C. The shank was brought into contact with the PEG-coated shuttle. PEG solidifies within seconds, forming a stable attachment. For assembly, both the shuttle and implant were held by hand, or the shuttle was held using tweezers. The flexible components of the implants should not be held by tweezers to avoid kinks or damage. Assembly was performed under direct visual observation and could be inspected afterwards under a microscope. In case of poor alignment or incomplete attachment, PEG was dissolved in water and the process was repeated.

### 2.6 Implantation

Implantation was evaluated in hydrogel and in ex vivo pancreas provided by a local butcher. The hydrogel was prepared by boiling 0.5 % w/v agarose (Agarose Standard ROTI Garose) in 150 mM phosphate-buffered saline (ROTI Cell PBS) and letting it cool in cuvettes. Both hydrogel and pancreas were at room temperature. The pancreas was not perfused.

The shuttle was held by tweezers and the assembly (with functional or dummy implants) was implanted into the hydrogel or pancreas to the full length of the shank. Carbon-tip tweezers allowed a more secure grip of the shuttle, whereas the shuttle slipped more easily when held by metal tweezers. In hydrogel, its transparency enabled visual observation and shuttles were removed after dissolution of PEG. In the pancreas, shuttles were removed after 5 min. Shuttles were removed using the same tweezers.

### 2.7 Electron microscopy and focused ion beam milling

Electron microscopy for surface analysis and imaging and focused ion beam (FIB) milling for the preparation of cross sections were performed with a ZEISS Auriga 40. The sample was sputtered with Au/Pd in the ratio 80/20. Images were acquired with a 3 kV acceleration voltage using the secondary electron detector with a stage tilt of 40.8° and no tilt correction.

## 3 Results and Discussion

### 3.1 Adaptations of the MEA to histological and functional conditions

Histology of the porcine pancreas showed a uniform distribution of islets but none within 250 µm of the organ’s surface (Figure 1). Therefore, our design focused on penetrating microelectrode arrays rather than placing the array on the surface of the pancreas. Penetrating MEAs include 2D arrays of micromachined needles with a single recording electrode at each tip, such as the Utah array or flexible devices (Steins *et al*., 2022). Other options integrate many electrodes along shanks, which can also be rigid (silicon-based) or flexible (polymer-based). In neuroscience, the “Butcher number” has been defined as the number of neurons destroyed per neuron recorded, with an estimate of 200 for Utah arrays and 2.5 for silicon shanks (Markus Meister, 2019)

**Figure 1:**
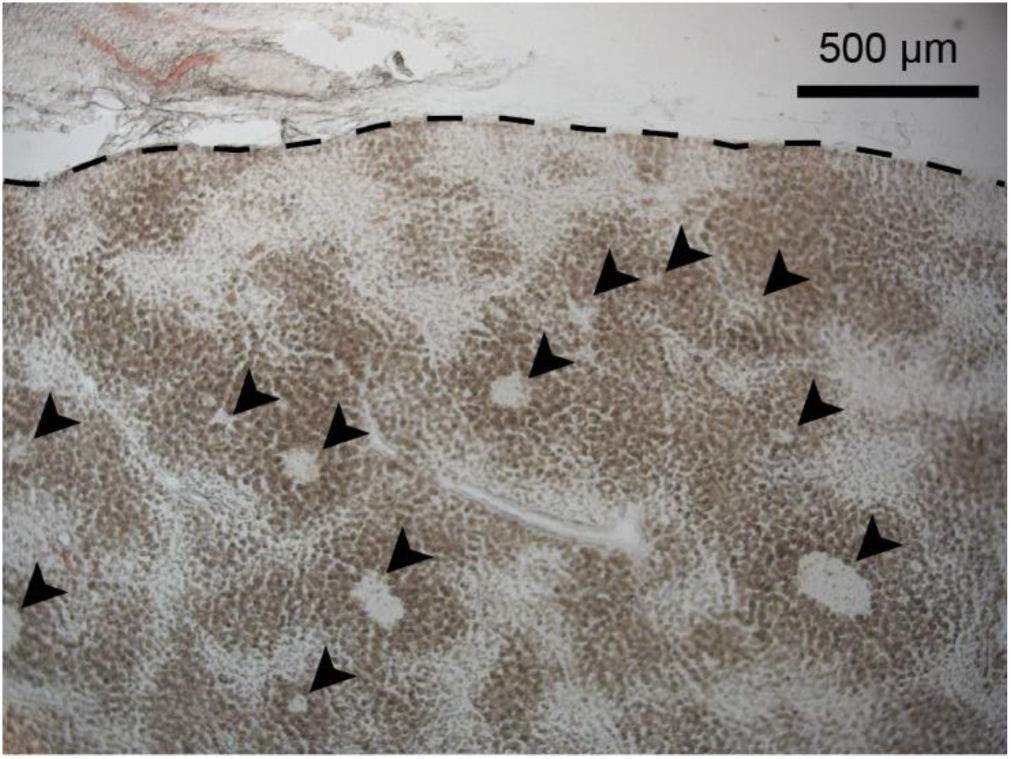
Semithin sliced porcine pancreas. Line indicates the pancreatic surface. Light spots marked by arrows indicate islets of Langerhans.

Flexible implants are preferable for chronic use (Du *et al*., 2017; Dai *et al*., 2018). Utah-array-like devices (even if flexible) would cause too much damage to the pancreas and are limited in depth compared to shank-based MEAs. Silicon or polymer shanks can be very long with electrodes placed along their length (Xu *et al*., 2021). The implantation of flexible MEAs is challenging but can be made possible by the use of insertion shuttles (Felix *et al*., 2013; Weltman, Yoo and Meng, 2016) . For neural applications, precise positioning is desirable but challenging. In the pancreas, positioning is less important. However, the highest density of islet cells is found in the tail of the pancreas (Ionescu-Tirgoviste *et al*., 2015), therefore implantation in this anatomic section may be optimal.

### 3.2 Design

The implant consists of a flexible polyimide-based MEA component connected to an EIB. The flexible MEA component (Figure 2) has a connector region, ribbon cable and shank. The 17 mm-wide connector region is designed for wire-bonding to the EIB with 66 bond pads at a pitch of 250 µm. The 4 cm-long ribbon cable is designed for simple handling. Eyelets near the shank support handling and fixation by gluing or suturing, which should stabilize the implant’s position in the organ to minimize movement artifacts.

**Figure 2:**
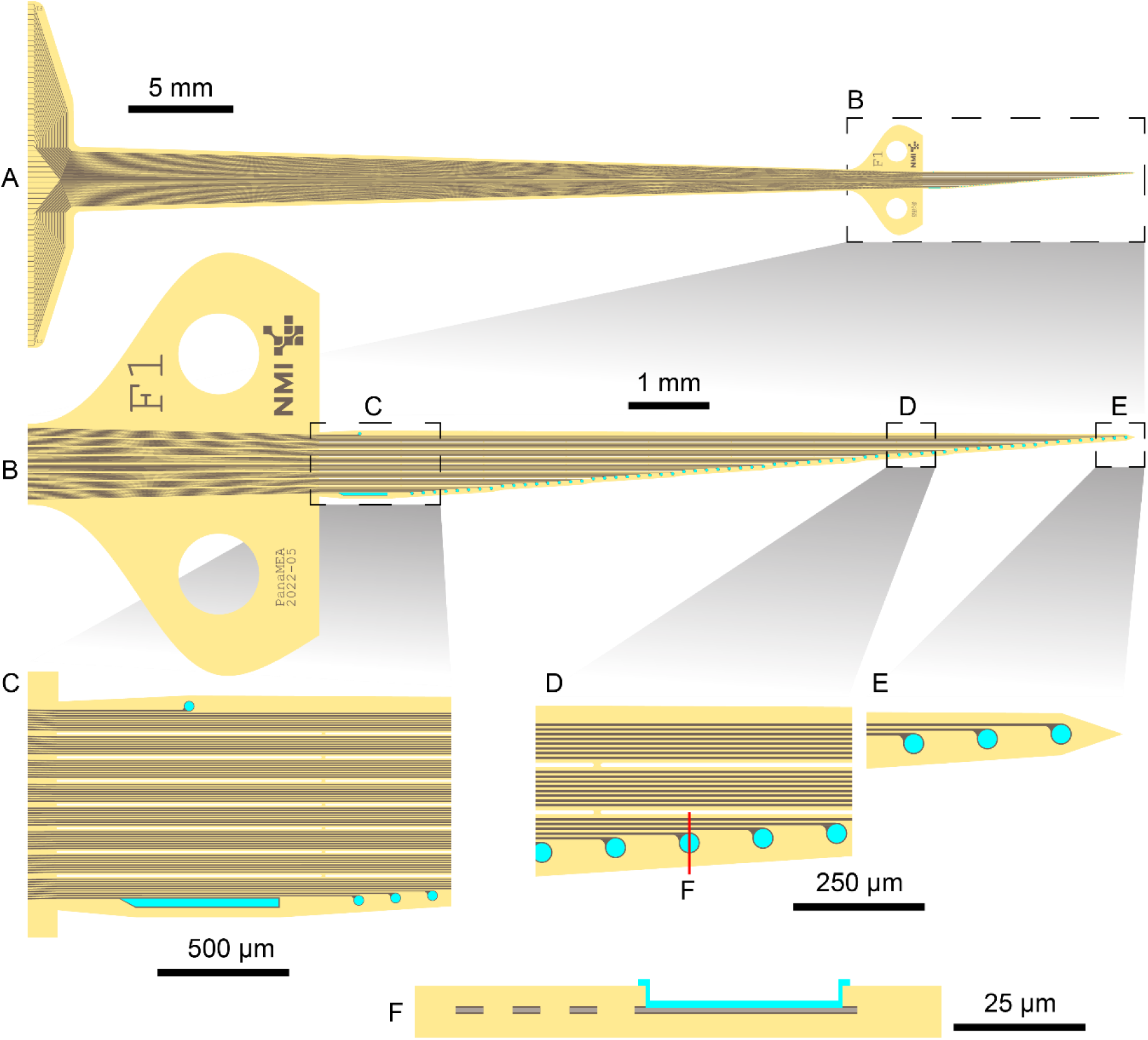
The design of the implant’s flexible microelectrode array component. Polyimide is indicated in yellow, insulated traces in brown and electrodes in teal. A: The MEA component is 5.5 cm long and includes a 17 mm-wide connector region, a ribbon cable, and a shank. B: Eyelets and shank. Eyelets include an identifier (here, F1). C: The proximal end of the shank includes a counter electrode (top) and reference microelectrode (bottom). D: Microelectrodes are distributed along one edge of the shank. Perforations (white) are included for every eight traces. E: The shank tapers to a sharp tip. F: A schematic cross-section of the red line in D through three traces and a microelectrode.

The 10 mm-long shank is intended for implantation into the pancreas (Figure 2B). A reference electrode and counter electrode are located proximally (Figure 2C), and 64 microelectrodes are distributed along one edge with a 140 µm pitch as the shank tapers to a sharp tip (Figure 2E). The microelectrodes and reference electrode have a diameter of 30 µm, and the rectangular counter electrode has an area of 17×10^3^ µm². For increased flexibility, the shank is perforated into eight filaments, each containing electrical traces for eight microelectrodes.

The shank’s maximum width of 850 µm is determined primarily by the resolution of the metal layer. Although we use contact lithography and have produced similar flexible electrode array components with traces of 2 µm (Albeck *et al*., 2022), we chose to have minimum trace width and spacing of 4.5 µm to ensure reasonable yields and reduce process complexity. Proximal from the shank, the trace width and spacing were increased to reduce the risk of fabrication defects. For more flexibility, the shank could be made 50% narrower by using 2 µm traces and spacing and its thickness could be reduced to 4 µm, however these changes would increase fabrication complexity and defect density.

The flexible component has a thickness of 12 µm for a maximum cross-section of 10200 µm², and its flexibility is increased by perforations. This should minimize injury and immune response during chronic implantations. Acute injury immediately after implantation will result primarily from the implant–shuttle assembly, due five-fold higher cross-section of the shuttle and PEG.

The overall dimensions allow seven components to be produced per 100 mm wafer. Longer components would greatly reduce this number, while shorter components would impede handling.

### 3.3 Fabrication and connectors

For laser-based rapid prototyping of dummies, we achieved minimum cut widths of 15 µm in polyimide or parylene C. Dummies were sufficient to study mechanical behavior despite a lower resolution than photolithography and the absence of metal layers. For functional implants, polyimide was used because it offers good biocompatibility and mechanical properties, suitability for microfabrication and insulation (Hassler, Boretius and Stieglitz, 2011) and years of stability in clinically approved flexible implants (Daschner *et al*., 2017).

A functional implant is shown in Figure 3, showing the flexibility of the polyimide component (Figure 3A) and the connection to the EIB (Figure 3B). High resolution structures match the expected design (Figure 3C, D). Optical inspection demonstrated a high yield of defect-free devices without broken or short-circuited traces. Further detail of the flexible MEA components was revealed by electron microscopy and preparation of a FIB cross-section. Figure 4A shows the eleven most distal microelectrode and the first perforation. Traces are faintly visible extending proximally from each microelectrode. The tip is magnified in Figure 4B, and Figure 4C shows a cross-section through a perforation, three traces and a microelectrode. The cross-section revealed that the actual trace width was ∼3 µm, reduced from a nominal width of 4.5 µm (Figure S1).

**Figure 3:**
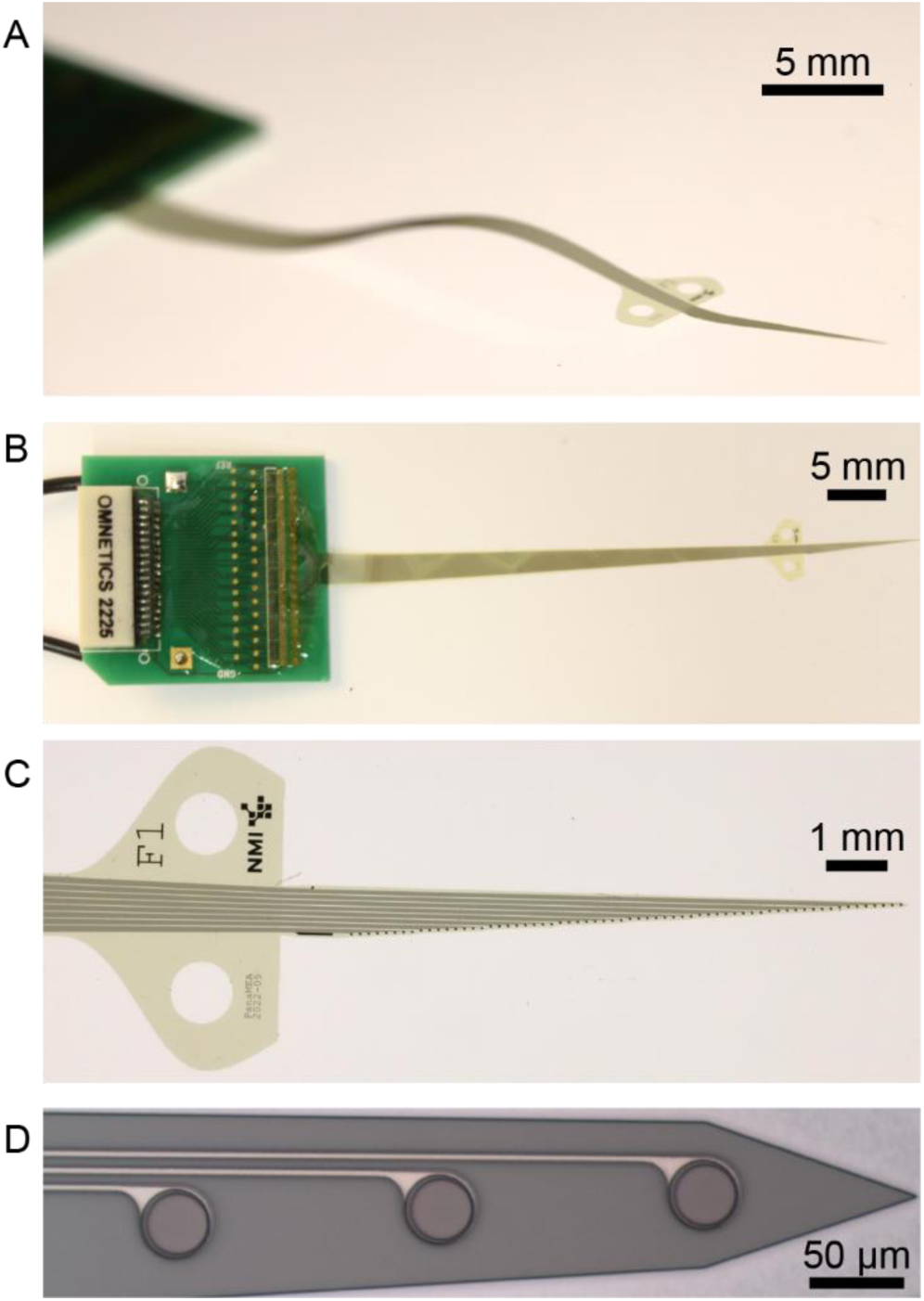
Optical images of the flexible implants. A: Photograph illustrating the flexibility of the implant. B: Top view of the implant including the EIB. C: Photograph of the shank and eyelets. D: Microscope image of the tip of the shank.

**Figure 4:**
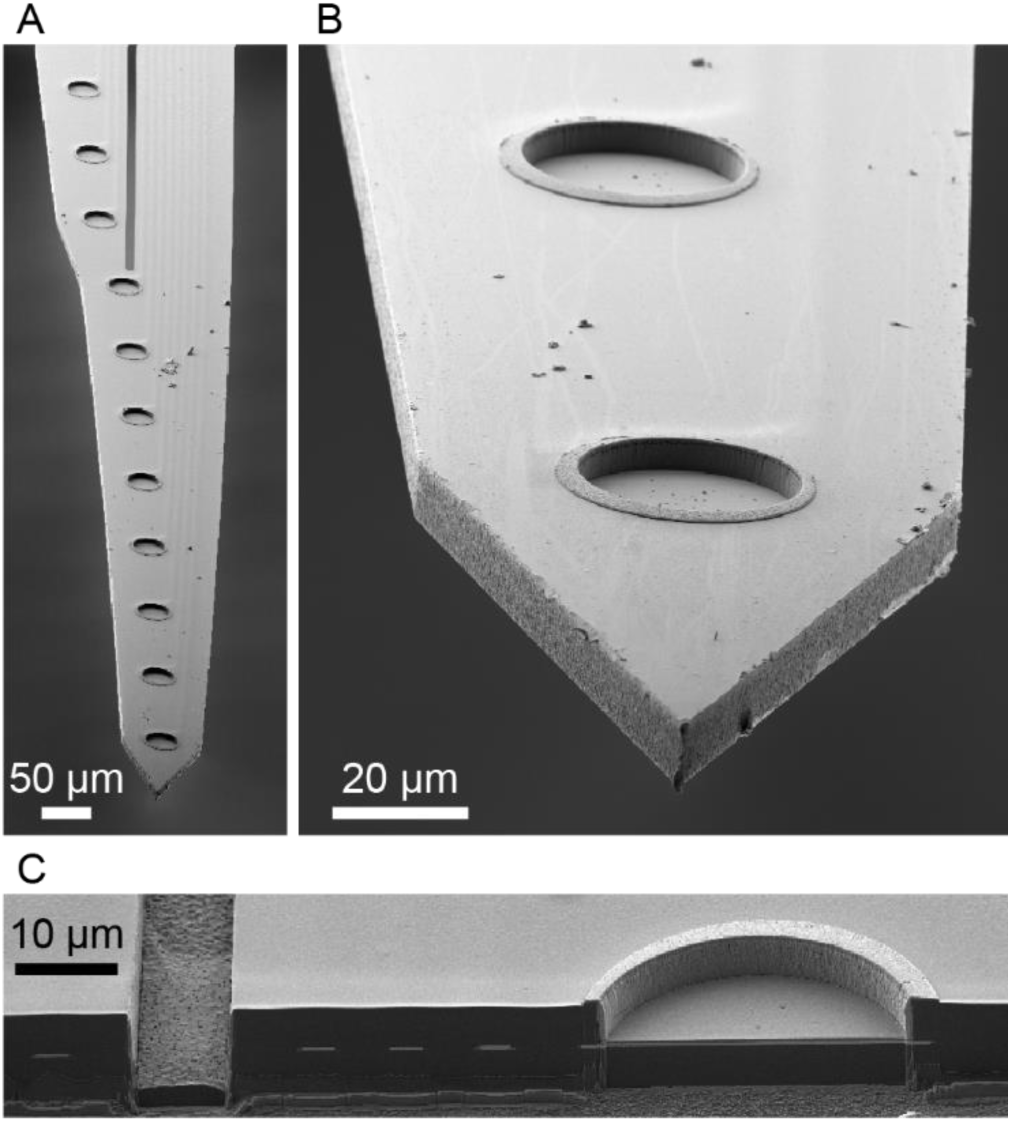
Electron microscopy. A: The tip of the shank including 11 microelectrodes and the first perforation. Traces are faintly visible. B: Magnification of the two distal electrodes and the tip of the implant. Electrodes are formed by etching the top polyimide layer to expose the traces, and patterning the recess with TiN. C: A cross-section prepared by FIB reveals the thin-film structures of the implant corresponding to the sketch in Figure 2F. A perforation (left) separates two polyimide filaments. Three traces are insulated by polyimide. The TiN electrode (right) is recessed into the polyimide. All images were obtained from the secondary electron (SE) detector.

The EIB includes two Omnetics Nano Strip connectors (A79024-001) soldered on opposite sides of the printed circuit board. These connectors were chosen due to their proven reliability ability in electrophysiology, reliability after a high number of mating cycles and their small size. Disadvantages include their cost (currently 65–85 € each) and the need for two well-aligned connectors on a single EIB to enable 64 channels. Low-cost single connectors with sufficient channels are available, such as Hirose DF40 connectors (∼2 € for 70 channels) (Voigts *et al*., 2020), however we decided against these due to the low number (<30) of mating cycles (Hirose Electric Co.,LTD, 2023).

### 3.4 Predicting the likelihood of recording beta cell activity

Previous studies have demonstrated in vitro recordings of beta cell activity, including the correlation of islet activity with glucose concentration (Pfeiffer *et al*., 2011) and recordings from the same islets over 30 days in vitro (Schönecker et al., 2014). We are unaware of any reports of the recording of islet activity in a large animal model, which has complications beyond these in vitro studies or small animal models.

We designed our implant to maximize the likelihood of recording the electrical activity of pancreatic beta cells while minimizing injury and the risk of pancreatitis. Signals of about 100 µV can be measured using microelectrodes in direct contact with human islets (Schönecker *et al*., 2015). Extracellular potentials decay with 1/*r* (Humphrey and Schmidt, 1990), suggesting that threshold detection at a five-sigma level with recording noise of 3 µVrms (Mierzejewski *et al*., 2020) may only be possible within a distance of ∼7.5 µm (Figure 5A). A proposed superficial implant (Harel, Tamar *et al*., 2001) would avoid penetrating trauma, but may not bring electrodes close enough to the pancreatic islets to record their activity.

**Figure 5:**
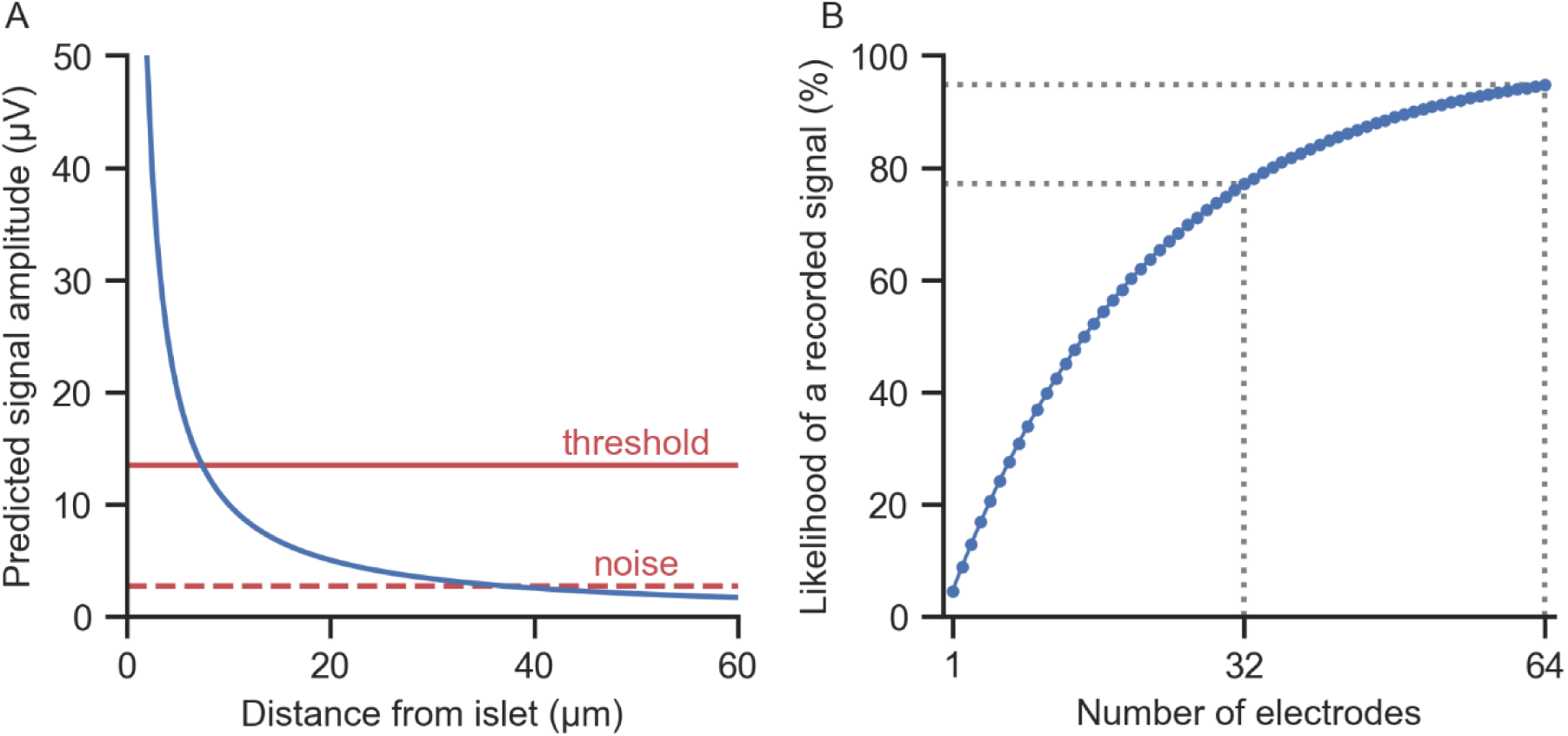
Left: Predicted signal amplitude vs. distance from an islet compared to expected recording noise of 2.7 µV and a five-sigma threshold. Right: Estimated likelihood of recording a signal with one electrode is 4.5 % and increases to 77 % and 95 % with 32 and 64 independently positioned microelectrodes, respectively.

We expect that a microelectrode placed randomly at a sufficient depth within the pancreas will have a certain likelihood of being near enough to an islet to record its activity, and that the likelihood of recording at least one useful signal should scale with the number of independent electrodes. Due to the rapid decay of extracellular signals with distance, we use the 4.5 % endocrine volume of the pancreas as the likelihood that a randomly placed microelectrode will be near enough to a beta cell to record its activity. If each microelectrode records from an independent volume, an array of *n* microelectrodes will therefore have a probability of *p_n_* = 1 – 0.955*^n^* of recording at least one useful signal. This probability is 77 % using 32 electrodes and 95 % for 64 electrodes (Figure 5). Importantly, this assumes a wide spacing between electrodes.

Factors related to the implantation and injury are not considered in this estimate and could reduce the likelihood of recording activity. Mechanical injury or effects of dissolved PEG could be expected to affect beta cell activity with acute or chronic consequences. Similar implants using similar methods have shown excellent results for chronic recordings in the brain (Du *et al*., 2017; Xu *et al*., 2017; Yang *et al*., 2019). In fact, most recordings with such implants are reported on dates after the implantation, after the animal has recovered from surgery. This raised the question of whether acute recordings were unsuccessful or were not attempted due the additional complexity of recording during surgery. Recently, acute recordings in the rat brain using a PEG-assisted implantation of flexible microelectrode arrays was reported (Xu *et al*., 2022). Still, the larger size of our implant–shuttle assembly or different injury mechanisms in the pancreas vs. the brain may affect the results of a chronic experiment.

Furthermore, the loss of islets during progression of diabetes mellitus type 2 must be considered. With an appropriate feedback loop for insulin administration, the goal would be that the loss of islets is slowed. Recording during progression of the disease could be informative, however the eventual loss of islets would prevent further benefit for the patient.

### 3.5 Electrochemical characterization

The electrical impedance of a microelectrode determines its intrinsic voltage noise (Mierzejewski *et al*., 2020). Representative impedance spectra are presented in Figure 6. Microelectrodes had an impedance magnitude at 1 kHz of 92±19 kΩ. The impedance spectra suggest that recordings using these electrodes in combination with a low-noise amplifier should have a noise as low as ∼3 µVrms, and that recording of extracellular potentials from beta cells is theoretically possible. Lower impedance and noise could be achieved by using PEDOT:PSS instead of TiN (Mierzejewski *et al*., 2020).

**Figure 6:**
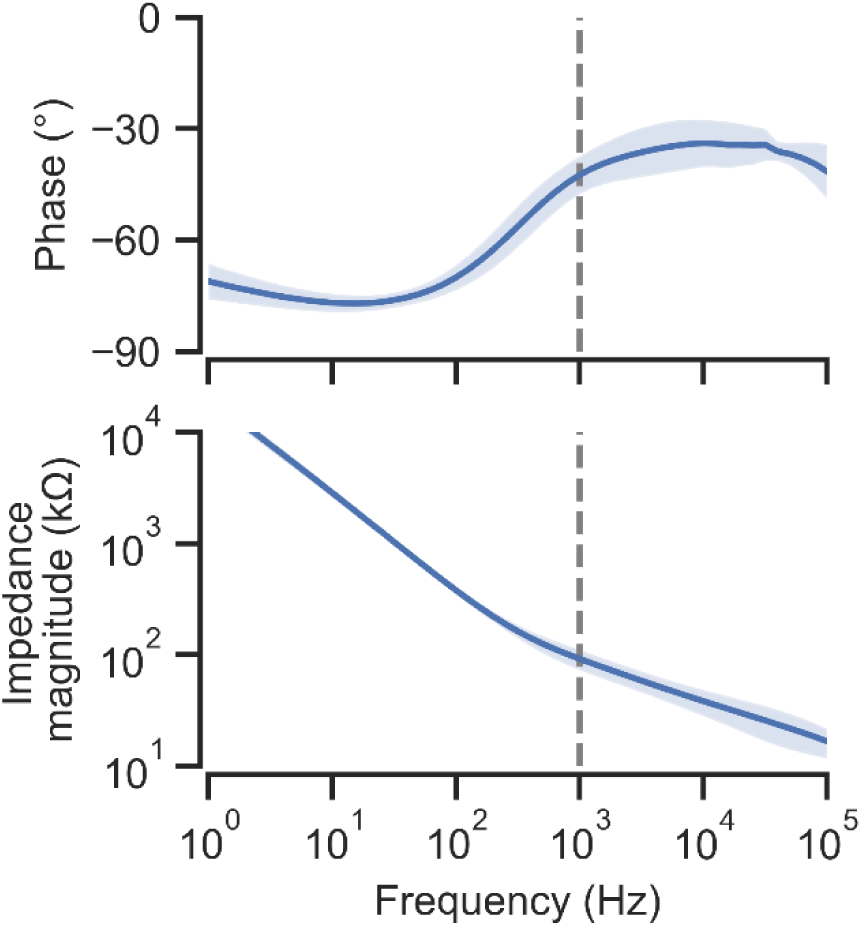
Bode plot of impedance spectra of TiN microelectrodes. Plots indicate mean and standard deviation for 64 electrodes. Impedance magnitude at 1 kHz was 92±19 kΩ (mean and standard deviation).

### 3.6 Implantation

Implantation of dummy implants in agarose hydrogel (Figure 7) revealed best results with tapered implants. In contrast, rectangular designs were more likely to become detached from the shuttle during implantations. Fully separated filaments (Guan *et al*., 2019) were tested but often became tangled which would prevent the intended spatial distribution of microelectrodes. Perpendicular supports at regular intervals ensured that the implanted dummies were untangled and fully extended.

**Figure 7:**
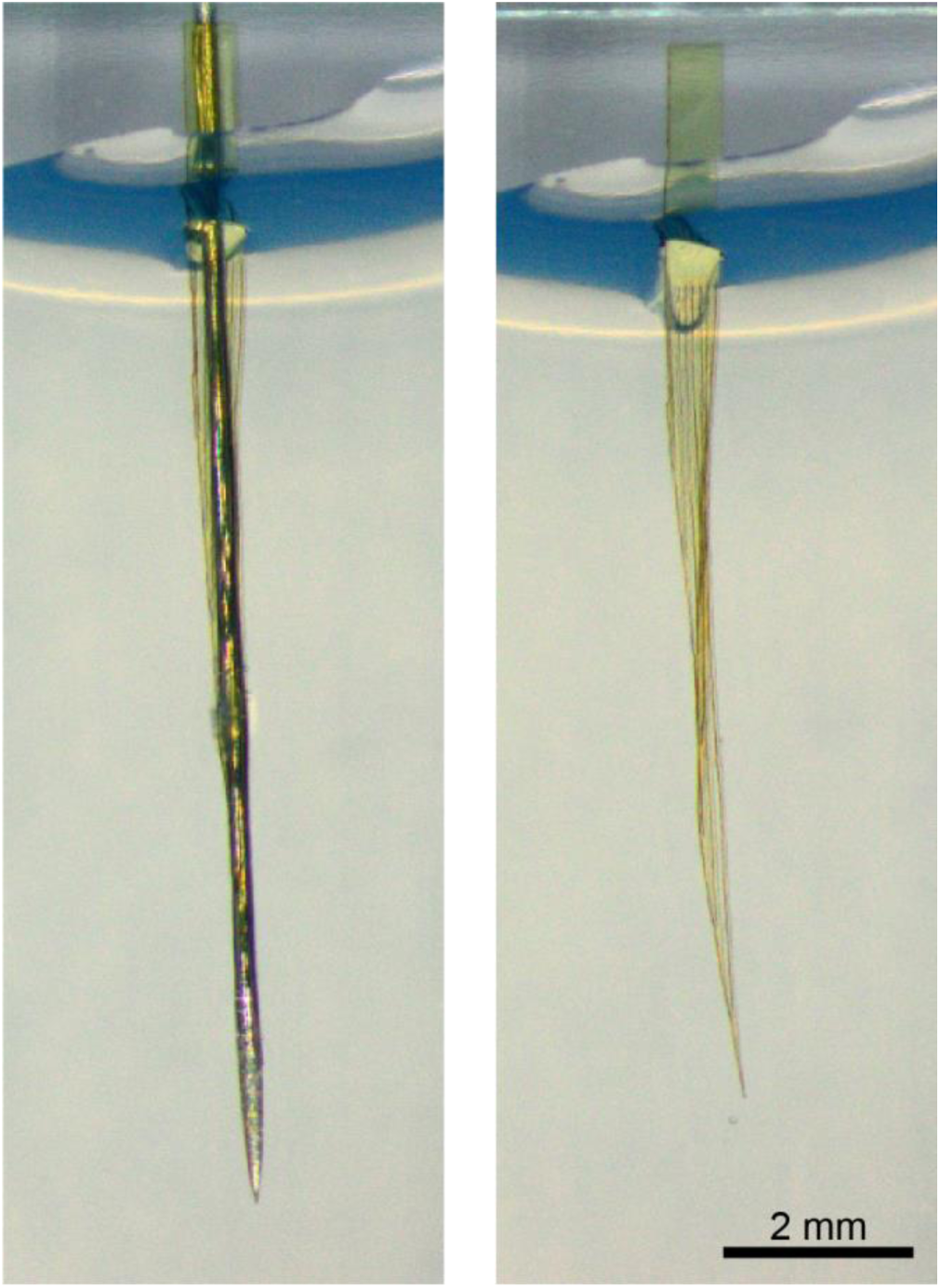
Implantation in agarose hydrogel. Left: a dummy implant–shuttle assembly was implanted 10 mm into the hydrogel. Right: Only the dummy implant remained after removal of the shuttle.

Uniform coating of the shuttle was achieved by manual dip-coating in PEG at a ∼45° angle. Proper coating avoided non-uniform droplet formation due to the Plateau–Rayleigh instability. The higher cross-section of droplets would make implantation more injurious, and the thicker PEG regions would require longer time to dissolve, delaying or preventing shuttle explantation. Any shuttles with such droplets should be recoated.

To allow separation of shuttle and implant, the entire PEG-coated region must be implanted. Only when in contact with fluid (in hydrogel or tissue) can the PEG dissolve. Therefore, shuttles must not be dipped in PEG too deeply. If necessary, saline can dissolve any PEG above the implanted depth.

Implantation in ex vivo pancreas demonstrated a high rate of success as well as two key failure modes (Figure 8). A total of 27 polyimide or parylene C dummies were implanted. Material-dependent differences were not observed. Dummy implantations were successful in 22/27 attempts (81%). In three cases (11%), the dummy implants detached from the shuttle, indicating too little PEG in the assembly. In two cases (7%), dummy implants were explanted with the shuttle, indicating that more time would have been needed to fully dissolve the PEG.

**Figure 8:**
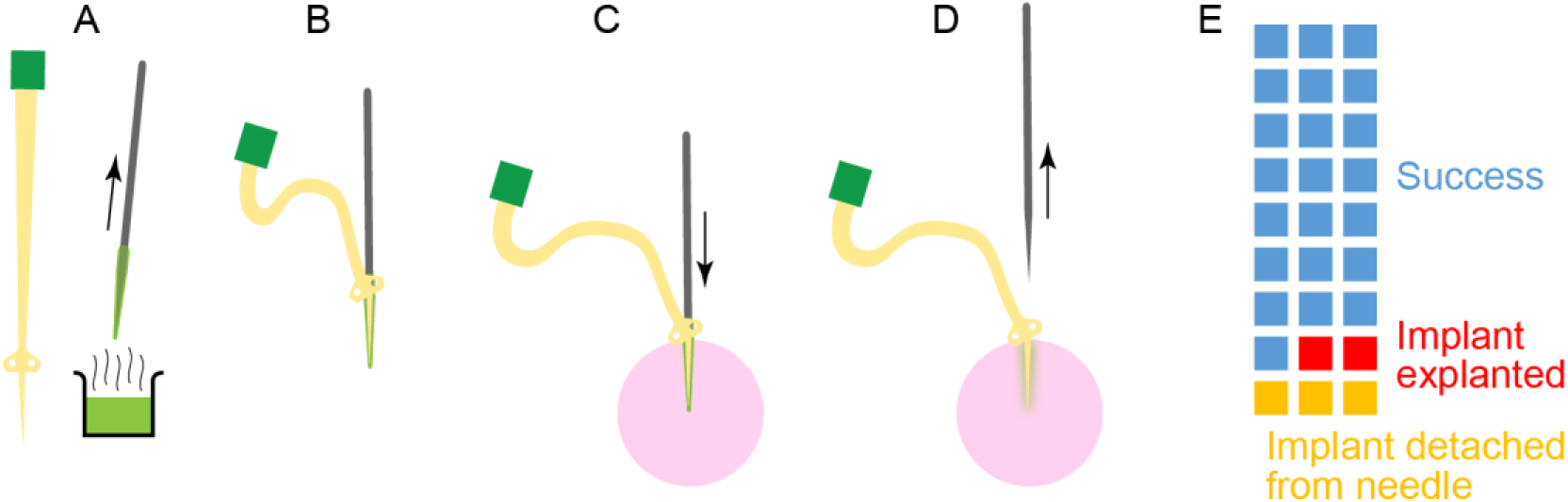
Implantation into ex vivo pancreas. Schematic illustration showing (A) dip-coating of the shuttle (grey) in hot PEG (green), (B) assembly of the implant–shuttle assembly, (C) implantation into the pancreas (pink), and (D) removal of the shuttle. E: Of 27 attempted implantations into ex vivo pancreas, 23 were successful (blue). Three implants detached from the shuttle during implantation (orange), indicating insufficient PEG to maintain adhesion. Two implants were explanted with the shuttle (red), indicating incomplete dissolution of PEG.

### 3.7 Suggestions for in vivo implantations

Care must be taken to use an appropriate amount of PEG to keep ensure adhesion during implantation, minimize the size of the implant–shuttle assembly and ensure reliable detachment and explanation of the shuttle. We prefer manual dip-coating and assembly (rather than an automatic process) due to fine handling and short time required before PEG solidifies. The amount and distribution of PEG should be measured for each assembly. Dissolution in water and recoating can easily correct assemblies with misalignment or excessive PEG.

The implant–shuttle assembly can be prepared before the surgery and should be stable for months under normal lab conditions. Assemblies should be protected from dust or contamination. Sterilization of PEG-based implant–shuttle assemblies has been reported with ethanol (Felix *et al*., 2013).

During in vivo recording we expect mechanical movements (e.g., peristalsis and respiration) may affect the placement of the implant. The eyelets (inner diameter of 1 mm) should allow fixation by suturing, however this remains to be evaluated. The eyelets also enable manipulation of the implant without touching the more sensitive electrical traces.

Our dual male Omnetics Nano Strip connectors (A79024-001) are compatible with amplifiers having female^1^ connectors (e.g. A79025-001) such as the 32-channel ME2100 from Multi Channel Systems MCS GmbH or 64-channel RHD headstages from Intan Technologies. We predicted a 77 % or 95 % chance of at least one successful signal with 32 or 64 channels, respectively. With multiple 64-channel implants, our prediction approaches 100 %. However, recordings in vivo may require connection to the amplifier before implantation. If at all possible, disconnecting and reconnecting implanted probes would require extreme care and would risk displacement or explantation.

The EIB and connectors here are appropriate for acute in vivo experiments. Chronic implants will require different connectors and encapsulation either for percutaneous or wireless telemetry.

While we have demonstrated successful implantation into ex vivo pancreas, the implantation into a living animal will have other challenges. Specifically related to the implantation, we expect that the force required for implantation may be higher. In comparison to an unperfused pancreas at room temperature, PEG may dissolve faster in the warm and perfused organ. Observation of the implantation may also be more challenging.

## 4 Conclusion and Outlook

We have designed an implant for intrapancreatic electrophysiology with an optimal likelihood of recording electrical activity of beta cells in the pancreatic islets in a large animal model. Characterization of the fabricated implants and demonstration of the shuttle-assisted implantation process suggest that the implants are suitable for their intended purpose.

In designing the implant, we have considered the physiology of the pancreas and the capabilities of flexible microelectrode arrays as developed for neural applications. Specifically, flexible, perforated implants should minimize mechanical injury and immune reactions for chronic implantation. As a next step, the performance of the implant should be evaluated in acute and later chronic animal experiments. While the size of the implant is small, the shuttle-based implantation will cause an acute injury, and effects on the beta cells or their electrical activity cannot be excluded. Ideally, activity of the beta cells would be controlled pharmacologically or by control of the animal’s blood glucose levels to prove the source of the recorded signals.

If animal studies can demonstrate reliable long-term recording, a medical device for treatment of diabetes mellitus could be envisioned. Such an implant could have a much higher lifetime than current enzyme-based biosensors. Connection to an insulin pump would enable more physiological insulin administration and potentially reduce symptoms and comorbidities of diabetes.

## 5 Acknowledgements

We thank Ilona Matiychyn (Multi Channel Systems MCS GmbH) for her expertise with PEG-assisted implantation of flexible devices and Helen Steins and Simon Werner for their expertise with assembly of the connectors.

## 6 Declarations

### 6.1 Ethical Approval

Not applicable

### 6.3 Authors’ contributions

D.P. and P.D.J. analyzed results and wrote the manuscript with input from all authors. M.T., A.K. and J.R.: histology. D.P., L.B., R.D., U.K., P.D.J.: conceived and designed the implants and implant–shuttle assembly. A.S.: device fabrication. R.D. and D.P.: implantation. D.P.: impedance measurements. M.K.: Electron microscopy and FIB. All authors reviewed the manuscript and approved its final version.

### 6.5 Avaiability of data and materials

Not applicable

---

**Figure S1:**
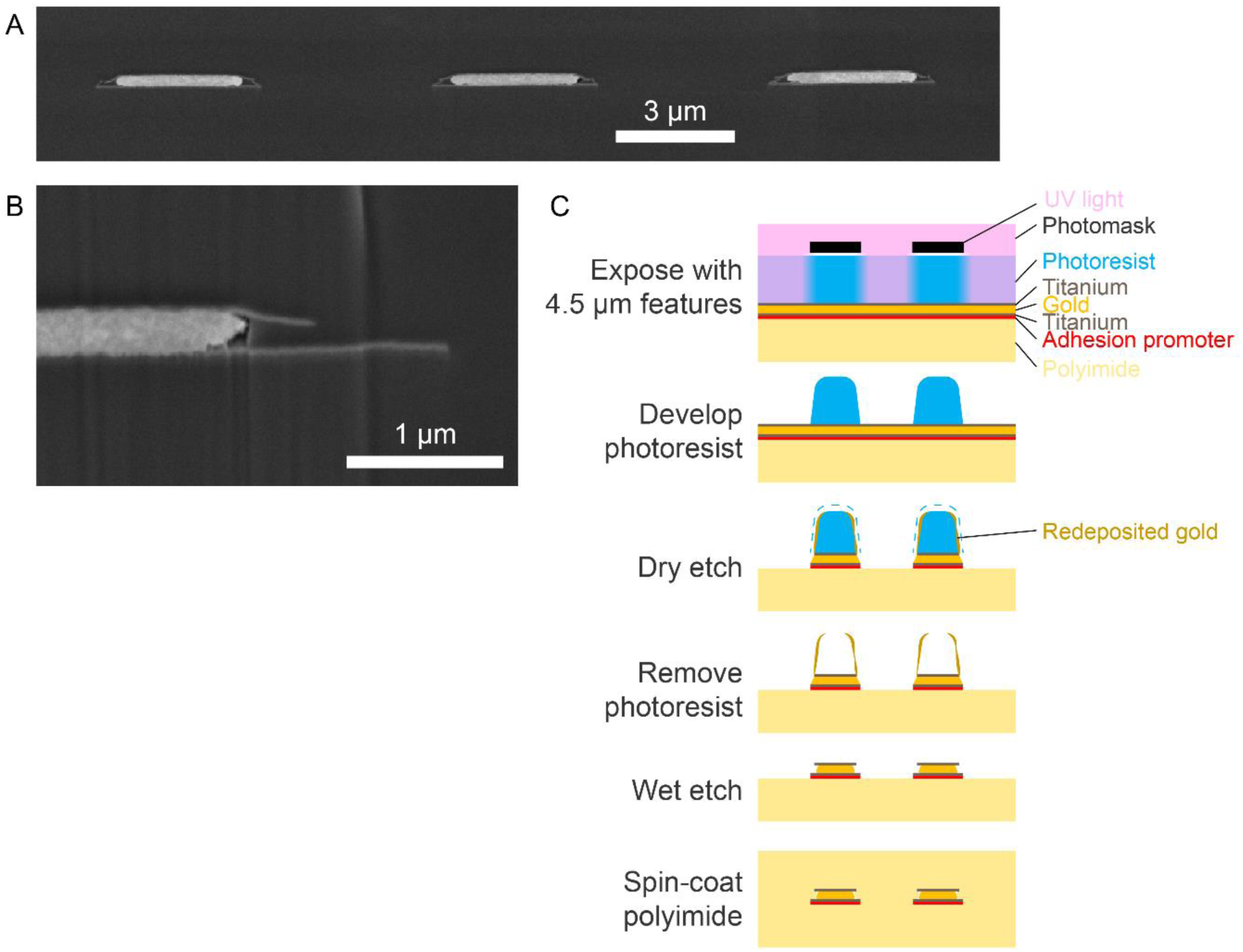
Thin film structure revealed by FIB-milled cross-section. A, B: Cross-section through traces shows underetching of gold between titanium adhesion layers. The upper titanium layer was free-standing where gold was underetched. A small void remained after spin-coating of the second polyimide layer. Images were obtained with the in-lens detector for elemental contrast. C: Schematic process illustrating the observed structures. Dry etching of gold causes redeposition on the photoresist. After photoresist removal, redeposited gold can cause short circuits. The redeposited gold can form long filaments, as it extends upon the entire length of the traces. A short dip in a gold etchant was sufficient to remove the redeposited gold, but also caused underetching of the gold between the titanium layers. The vertical scale of the illustration is modified for clarity. Photoresist was 3 µm thick. Each polyimide layer was 6 µm thick. The metal thickness was 0.3 µm. The adhesion promoter is too thin to be apparent in the SEM images.

## Footnotes

1 The apparent gender of Omnetics Nano Strip connectors is misleading.

